# Association of Sleep with Anxiety in Chinese Empty-Nest Older Adults: The Mediating Roles of Physical Exercise and Social Participation

**DOI:** 10.64898/2026.08.06.743184

**Authors:** Yiming Ma, Quan Zhou, Jiaran Jiang, Naizhuo Zhang

## Abstract

Sleep disturbances are common among older adults and are often closely associated with anxiety symptoms, which together can substantially compromise physical and mental health.A cross-sectional analysis was conducted using data from the 2017/2018 wave of the Chinese Longitudinal Healthy Longevity Survey (CLHLS). The study included 6,106 Chinese older adults (aged ≥65 years) living in empty-nest households. Physical exercise was measured based on self-reported regular engagement in exercise, and social participation was defined as involvement in organized social activities (e.g., community or group-based events). Associations between sleep quality and duration and anxiety were estimated using multivariable logistic regression, and mediation effects of physical exercise and social participation were tested using bootstrap resampling.Among the 6,106 participants, 593 (9.7%) reported anxiety symptoms. Multivariable logistic regression analysis indicated that poor sleep quality was significantly associated with a higher risk of anxiety (OR = 3.23, 95% CI: 2.62–4.01, P < 0.001). Short sleep duration was also an independent risk factor for anxiety (OR = 1.34, 95% CI: 1.04–1.74, P = 0.025), whereas long sleep duration showed no significant association. Subgroup analyses stratified by sex, marital status, and economic status consistently revealed significant associations between sleep (both quality and duration) and anxiety, with no statistically significant interactions observed. Mediation analyses demonstrated that physical exercise significantly mediated the relationships of both sleep quality and long sleep duration with anxiety (all P < 0.01, with bootstrap confidence intervals excluding zero). Similarly, social participation significantly mediated the relationships of both short and long sleep duration with anxiety (all P < 0.01, with bootstrap confidence intervals excluding zero).In Chinese empty-nest older adults, both poor sleep quality and short sleep duration are independent risk factors for anxiety symptoms, with sleep quality exerting a stronger influence. Physical exercise partially mediated the associations between sleep quality and anxiety and between long sleep duration and anxiety, whereas social participation mediated the association between sleep duration (short and long) and anxiety. Improving sleep quality and ensuring adequate sleep duration may help reduce anxiety risk, and interventions promoting physical exercise and social participation could enhance these protective effects.

## Introduction

With rapid population aging, the number of empty-nest older adults in China has been increasing markedly. Empty-nest older adults are defined as older individuals whose children are absent from the household and therefore unable to provide day-to-day support; they typically live alone or only with a spouse ^[1]^. According to the Chinese Longitudinal Aging Society Survey, empty-nest older adults accounted for 47.5% of China’s older population in 2016, increased to 59.7% by 2021, and are projected to reach 90% by 2030^[2][3]^. This steep increase and the anticipated high future prevalence constituting a major public health and social challenge in the context of population aging in China. Relevant research shows that the empty-nest status has adverse effects on the physical health, cognitive function, and mental health of older adults^[4]^. Some even develop “empty-nest syndrome,” characterized by sadness, feelings of loss, helplessness, and difficulties adapting to changes in family roles^[5][6]^, which in severe cases may progress to depression or anxiety^[7]^.Other researchers pointed out that for empty-nest syndrome, emotional and behavioral symptoms may diminish or disappear over time^[8]^. The physical health and psychological problems of empty-nest older adults in China require urgent attention and intervention.

Sleep disturbances are highly prevalent among older adults. With advancing age, sleep duration gradually shortens and the incidence of sleep disturbances increases significantly^[9]^. Poor sleep is associated with various adverse health outcomes, including diminished physical function, cognitive impairment, and reduced social engagement. Meta-analyses of randomized controlled trials indicate that improving sleep quality can significantly benefit mental health^[10]^. Epidemiological studies have shown that both poor sleep quality and insufficient sleep duration markedly elevate the risk of anxiety and depression^[11]^. Among Chinese older adult population, sleep duration is negatively correlated with anxiety—shorter sleep duration is associated with a higher risk of anxiety^[12]^.Although substantial evidence highlights the critical role of sleep in older adults’ mental health, the comprehensive underlying mechanisms remain insufficiently understood. In order to fill the research gap, this study aims to elucidate the intrinsic mechanisms by which sleep quality influences anxiety, specifically by examining the mediating pathways of physical exercise and social participation.

Anxiety represents a prevalent psychological concern among older adults. Anxiety disorders encompass conditions such as separation anxiety disorder, selective mutism, specific phobia, social anxiety disorder, generalized anxiety disorder, panic disorder, and agoraphobia. Their high prevalence, chronicity, and frequent comorbidity have led the World Health Organization to rank anxiety disorders as the ninth leading cause of health-related disability globally^[13]^. Researcher has established a bidirectional relationship between sleep and anxiety: sleep disturbances can heighten the risk of anxiety, which, in turn, can contribute to a deterioration in sleep quality^[14]^. Investigating the mechanisms underlying this relationship and identifying relevant intervention pathways are significant importance for enhancing the quality of life among older adults.

Physical exercise demonstrates significant benefits in improving sleep and alleviating anxiety among older adults. For instance, a 12-week Tai Chi program significantly improved sleep quality, reduced perceived pain, and death anxiety in older women (both P < 0.01), highlighting the value of traditional exercise for this population^[15]^. Modern exercise interventions similarly show positive effects^[16]^. Furthermore, systematic reviews confirmed that exercise can improve sleep and ameliorate symptoms of anxiety and depression^[17]^.Higher levels of social participation are also associated with better mental health, quality of life, and cognitive function in older adults, along with a reduced risk of anxiety and depression. Among older adults with multimorbidity, social participation is closely connected to sleep quality^[18]^.Research indicated that social participation may influence insomnia indirectly through sequential mediators such as frailty, anxiety, and depression. Thus, social participation might alleviate anxiety and subsequently improve sleep quality/duration^[19]^. Collectively, this evidence suggests that physical exercise and social participation may serve as mediators in the relationship between sleep and anxiety.

In summary, although existing literature suggests interconnections among sleep, anxiety, physical exercise, and social participation in the mental health of older adults, research specifically focusing on the empty-nest population remains limited, particularly with respect to underlying mediating mechanisms. Accordingly, this study examines the associations between sleep quality and sleep duration and anxiety among Chinese empty-nest older adults, and further investigates the mediating roles of physical exercise and social participation. The findings aim to provide empirical evidence to inform targeted interventions to improve mental health and quality of life in this vulnerable population.

## Methods

### Data Source and Participants

The Chinese Longitudinal Healthy Longevity Survey (CLHLS) is a nationally representative prospective cohort study designed to investigate factors associated with health and longevity. It is conducted by the Center for Healthy Aging and Development Studies and the National School of Development at Peking University, covering 23 provinces, municipalities, and autonomous regions across China. The survey targets individuals aged 65 years or older and their adult children (aged 35–64 years). The survey was initiated in 1998, with follow-up waves conducted in 2000, 2002, 2005, 2008–2009, 2011–2012, 2014, and 2017–2018. The study protocol was approved by the Biomedical Ethics Review Committee of Peking University (IRB00001052-13074). To ensure data timeliness, the present analysis focused on the 2017–2018 wave, which included 15,874 participants. After applying the following exclusion criteria, the final analytical sample comprised 6,106 participants: (1) aged <65 years (n=95); (2) missing data on key variables (sleep quality or sleep duration, n=2,760); (3) missing covariate information (n=2,270); and (4) not classified as empty-nest older adults (n=4,643).

### Anxiety

Anxiety was assessed using the 7-item Generalized Anxiety Disorder scale (GAD-7) included in the CLHLS survey, which evaluates the frequency of anxiety symptoms over the past two weeks^[20]^. The GAD-7 has demonstrated excellent internal consistency (Cronbach’s α = 0.92) and good test-retest reliability (intraclass correlation coefficient [ICC] = 0.83). The scale consists of seven items, each scored on a 4-point Likert scale: 0 (not at all), 1 (several days), 2 (more than half the days), and 3 (nearly every day). Summed scores range from 0 to 21, with higher scores indicating greater anxiety severity. Consistent with prior studies, a total score ≥5 was used to indicate the presence of anxiety symptoms.

### Sleep Quality and Sleep Duration

Sleep quality and sleep duration were assessed using two self-reported questions: “How do you rate your current sleep quality?” and “How many hours of sleep do you get on average per day?”, both of which are widely used in epidemiological studies.^[21][22]^. Sleep quality was originally rated on a 5-point scale: excellent, good, fair, poor, or very poor. Consistent with the approach used by Gu et al.^[23]^, responses were dichotomized into “good sleep quality” (excellent or good) and “poor sleep quality” (fair, poor, or very poor). Sleep duration was classified according to the recommendations of the National Sleep Foundation for older adults (7–8 hours per night). Participants were categorized into three groups: short sleep duration (<7 hours), normal sleep duration (7–8 hours), and long sleep duration (>8 hours).^[24]^

### Physical Exercise and Social Participation

Physical exercise was assessed using the question: “Do you regularly engage in physical exercise (referring to purposeful fitness activities such as walking, playing ball, running, or qigong)?” Responses were dichotomized as yes (coded as 1) or no (coded as 0). Social participation was evaluated with the question: “Do you participate in any organized social activities?” Original responses were on a 5-point ordinal scale ranging from “1, almost every day” to “5, never.” For analysis, this variable was dichotomized: participants who selected “never” were classified as non-participants (coded as 1), and all others were classified as participants (coded as 0). In this study, “social activities” specifically refer to organized activities, thereby capturing a proactive form of social engagement distinct from routine daily tasks such as housework, gardening, or caring for livestock.

### Covariates

Covariates encompassed demographic characteristics, socioeconomic status, and lifestyle factors. Demographic variables included age (in years), sex (male/female), and marital status, which was dichotomized as “married and living with spouse” versus “other” (including separated, divorced, widowed, or never married). Socioeconomic indicators included educational attainment (no formal education vs. more than one year of schooling), residential area (urban, town, or rural), and self-rated financial status (affluent or above vs. average or below). Lifestyle factors comprised smoking status (current smoker: yes/no) and drinking status (current drinker: yes/no). Participants were classified as current smokers or current drinkers if they reported any present use, regardless of frequency or amount.

### Statistical Analysis

All statistical analyses were conducted using R software (version 4.4.2). Continuous variables were tested for normality. Normally distributed continuous variables are presented as mean ± standard deviation (SD), with group comparisons conducted using one-way analysis of variance (ANOVA). Non-normally distributed continuous variables are presented as median (interquartile range, IQR). Categorical variables were presented as frequencies and percentages, and group differences were assessed using the chi-square test. Multivariable binary logistic regression models estimated odds ratios (ORs) with 95% confidence intervals (CIs) for associations between sleep quality and sleep duration and anxiety symptoms. To control for potential confounders, three sequentially adjusted models were fitted: a crude model (Model 1); Model 2 adjusted for sex, age, and marital status; and Model 3 additionally adjusted for educational attainment, residential area, financial status, smoking, and drinking status. Subgroup analyses were performed to examine potential effect modification by sex, age, and financial status on the primary sleep-anxiety associations. Finally, mediation analyses evaluated the indirect effects of physical exercise and social participation on the relationships between sleep (quality and duration) and anxiety. Direct, indirect, and total effects were estimated using the nonparametric bootstrap method with 5,000 resamples, and 95% biascorrected CIs were derived. A mediation effect was considered statistically significant if the 95% CI for the indirect effect did not include zero.

## Results

### Baseline Characteristics

A total of 6,106 older adults were included in this study. Using a GAD-7 cutoff score of ≥5, participants were categorized into an anxiety group (n = 593, 9.7%) and a non-anxiety group (n = 5,513, 90.3%). The overall mean age was 79.3 years (SD = 9.7), with no significant difference between the two groups (p = 0.908). Significant differences in sociodemographic characteristics were observed between the groups. Compared with the non-anxiety group, the anxiety group had a lower proportion of urban residents (16.7% vs. 24.9%) and a higher proportion of rural residents (46.4% vs. 42.2%). The anxiety group also contained a significantly higher percentage of females (56.7% vs. 45.0%) and individuals reporting an average or below financial status (90.4% vs. 80.3%). Moreover, the illiteracy rate was higher in the anxiety group (46.2% vs. 34.9%). Health-related behaviors also differed significantly. The anxiety group had a markedly higher prevalence of short sleep duration (71.7% vs. 54.2%) and a lower prevalence of long sleep duration (14.0% vs. 22.6%). Poor sleep quality was reported by 76.1% of the anxiety group, compared with 44.7% in the non-anxiety group. Additionally, regular physical exercise (30.7% vs. 39.3%) and social participation (14.0% vs. 19.1%) were less common in the anxiety group.

### Association of Sleep Quality/Duration with Anxiety

Multivariable logistic regression analyses examined the associations between sleep quality, sleep duration, and anxiety (Table 2). In the unadjusted model (Model 1), poorer sleep quality was significantly associated with a higher likelihood of anxiety (OR = 3.55, 95% CI: 2.88–4.39, p < 0.001). This association remained significant after adjusting for age, sex, and marital status (Model 2; OR = 3.45, 95% CI: 2.80–4.27, p < 0.001) and after further adjustment for residential area, financial status, educational attainment, smoking, and drinking (Model 3; OR = 3.23, 95% CI: 2.62–4.01, p < 0.001). Although the strength of the association showed a stepwise attenuation across models, it remained consistently significant, indicating a robust relationship between poor sleep quality and anxiety. Regarding sleep duration, short sleep duration was significantly associated with anxiety in the unadjusted model (Model 1; OR = 1.36, 95% CI: 1.06–1.76, p = 0.018). The association persisted after sequential adjustment for demographic factors (Model 2; OR = 1.33, 95% CI: 1.04–1.72, p = 0.028) and additional socioeconomic and lifestyle factors (Model 3; OR = 1.34, 95% CI: 1.04–1.74, p = 0.025), demonstrating relative stability across models.

**Table 1.** Baseline Characteristics of Participants by Anxiety Status.

|  | [ALL] N=6106 | Non-anxious N=5513 | Anxious N=593 | p.overall |
| --- | --- | --- | --- | --- |
| **Age Mean (SD)** | 79.252 (9.690) | 79.249 (9.680) | 79.272 (9.788) | 0.908 |
| **Residence, N (%):** |  |  |  | <0.001 |
| **City** | 1470 (24.075%) | 1371 (24.868%) | 99 (16.695%) |  |
| **Town** | 2034 (33.311%) | 1815 (32.922%) | 219 (36.931%) |  |
| **Rural** | 2602 (42.614%) | 2327 (42.209%) | 275 (46.374%) |  |
| **Sex, N (%):** |  |  |  | <0.001 |
| **Female** | 2817 (46.135%) | 2481 (45.003%) | 336 (56.661%) |  |
| **Male** | 3289 (53.865%) | 3032 (54.997%) | 257 (43.339%) |  |
| **Marital, N (%):** |  |  |  | 0.010 |
| **Married and living with partner** | 1764 (28.890%) | 1565 (28.387%) | 199 (33.558%) |  |
| **Others** | 4342 (71.110%) | 3948 (71.613%) | 394 (66.442%) |  |
| **Economic status, N (%):** |  |  |  | <0.001 |
| **Average and below** | 4965 (81.313%) | 4429 (80.337%) | 536 (90.388%) |  |
| **Affluent and above** | 1141 (18.687%) | 1084 (19.663%) | 57 (9.612%) |  |
| **Sleep duration, N (%):** |  |  |  | <0.001 |
| **Short sleep duration** | 3411 (55.863%) | 2986 (54.163%) | 425 (71.669%) |  |
| **Normal sleep duration** | 1367 (22.388%) | 1282 (23.254%) | 85 (14.334%) |  |
| **Long sleep duration** | 1328 (21.749%) | 1245 (22.583%) | 83 (13.997%) |  |
| **Smoke, N (%):** |  |  |  | 0.023 |
| **No** | 4974 (81.461%) | 4470 (81.081%) | 504 (84.992%) |  |
| **Yes** | 1132 (18.539%) | 1043 (18.919%) | 89 (15.008%) |  |
| **Drink, N (%):** |  |  |  | 0.004 |
| **No** | 5002 (81.919%) | 4490 (81.444%) | 512 (86.341%) |  |
| **Yes** | 1104 (18.081%) | 1023 (18.556%) | 81 (13.659%) |  |
| **Exercise, N (%):** |  |  |  | <0.001 |
| **Yes** | 2350 (38.487%) | 2168 (39.325%) | 182 (30.691%) |  |
| **No** | 3756 (61.513%) | 3345 (60.675%) | 411 (69.309%) |  |
| **Education, N (%):** |  |  |  | <0.001 |
| **Illiterate** | 2197 (35.981%) | 1923 (34.881%) | 274 (46.206%) |  |
| **More than one year of education** | 3909 (64.019%) | 3590 (65.119%) | 319 (53.794%) |  |
| **Sleep quality, N (%):** |  |  |  | <0.001 |
| **Good sleep quality** | 3191 (52.260%) | 3049 (55.306%) | 142 (23.946%) |  |
| **Poor sleep quality** | 2915 (47.740%) | 2464 (44.694%) | 451 (76.054%) |  |
| **Social participation, N (%):** |  |  |  | 0.003 |
| **Yes** | 1135 (18.588%) | 1052 (19.082%) | 83 (13.997%) |  |
| **No** | 4971 (81.412%) | 4461 (80.918%) | 510 (86.003%) |  |

**Table 2.**
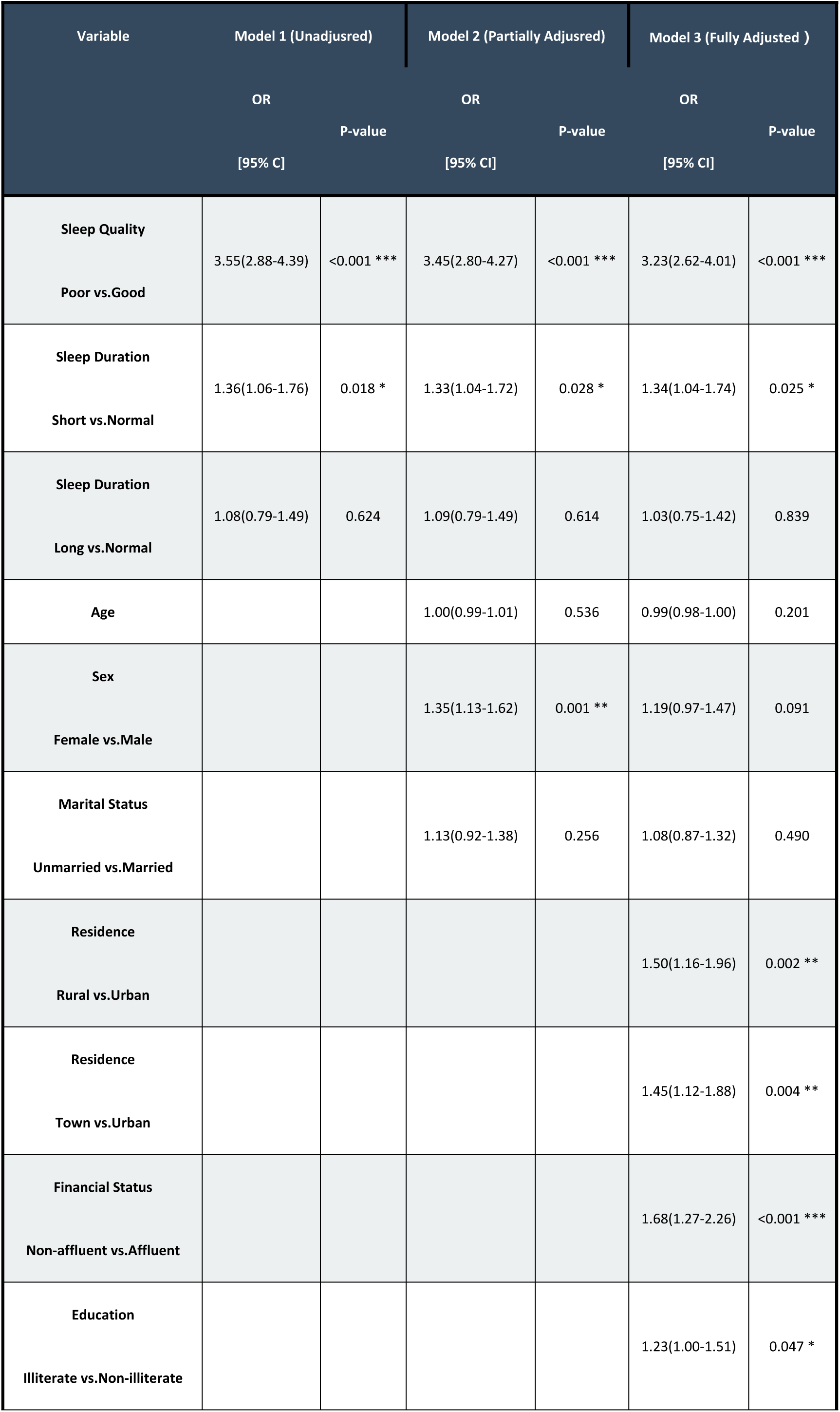
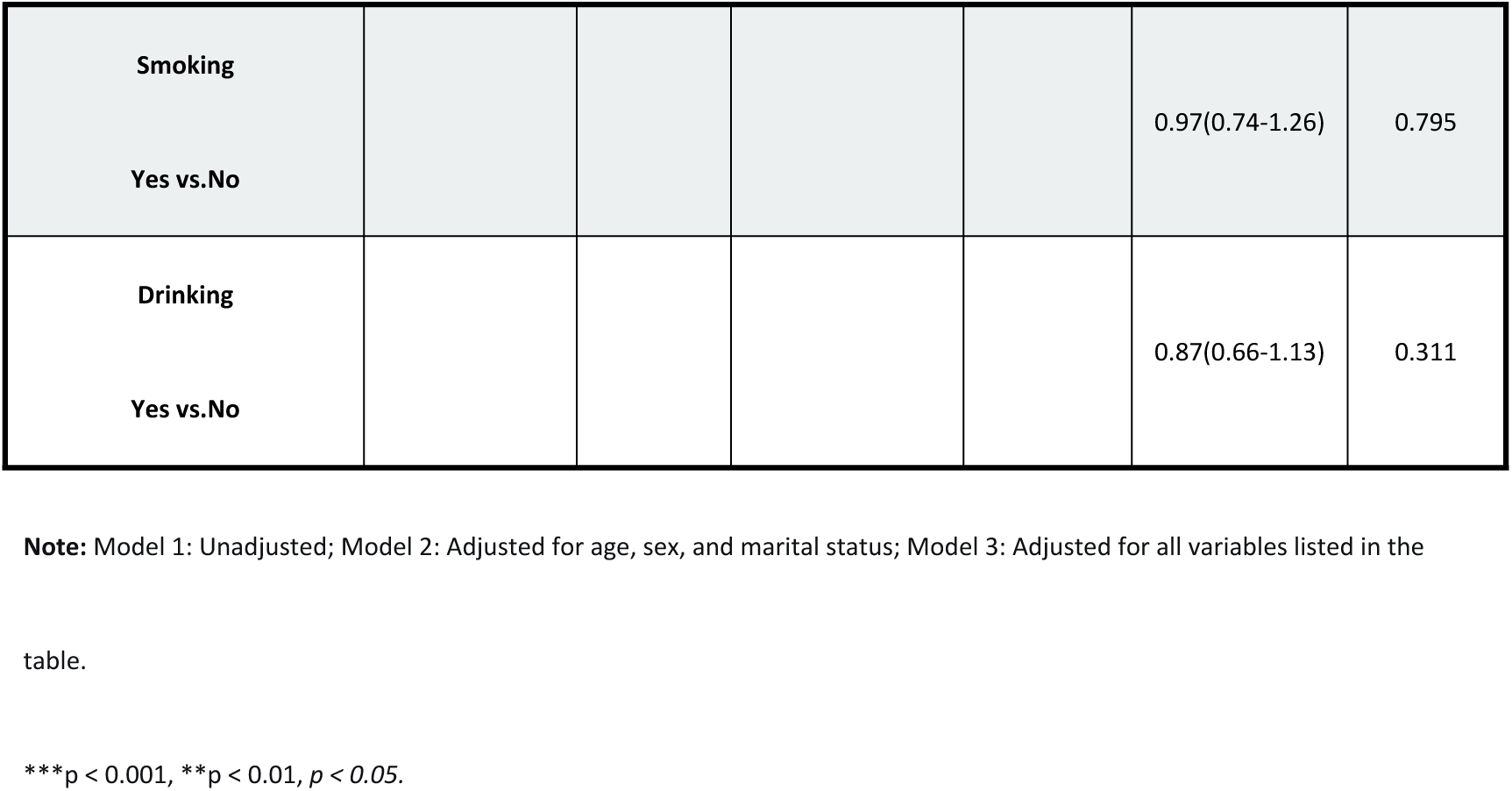
Multivariate Logistic Regression Analysis of the Association of Sleep Quality and Sleep Duration with Anxiety.

### Association of Sleep Quality and Sleep Duration with Anxiety in Different Subgroups

Subgroup analyses examined the consistency of associations between sleep quality, sleep duration, and anxiety across strata defined by sex, marital status, financial status, smoking status, drinking status, educational attainment, and residential area. No significant effect modification was observed for any of these covariates (all p or interaction > 0.05; see Figure 2A, B). Of note, the association between poor sleep quality and anxiety showed borderline significance for interaction with marital status (p for interaction = 0.07; Figure 2A). Short sleep duration remained significantly associated with higher odds of anxiety (p < 0.001; Figure 2B), whereas no significant association was observed for long sleep duration (p = 0.737; Figure 2B).

**Fig. 1.**
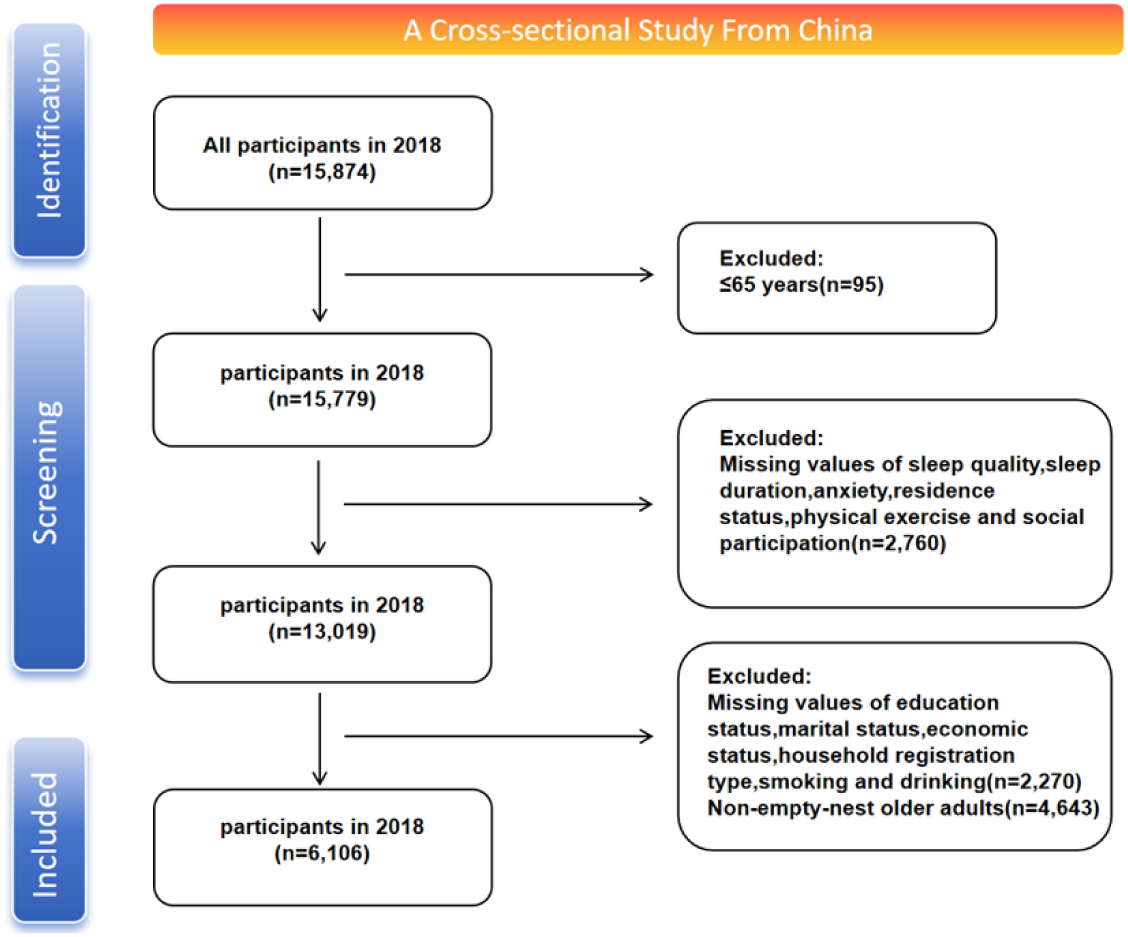
Sample selection flowchart

**Fig. 2.**
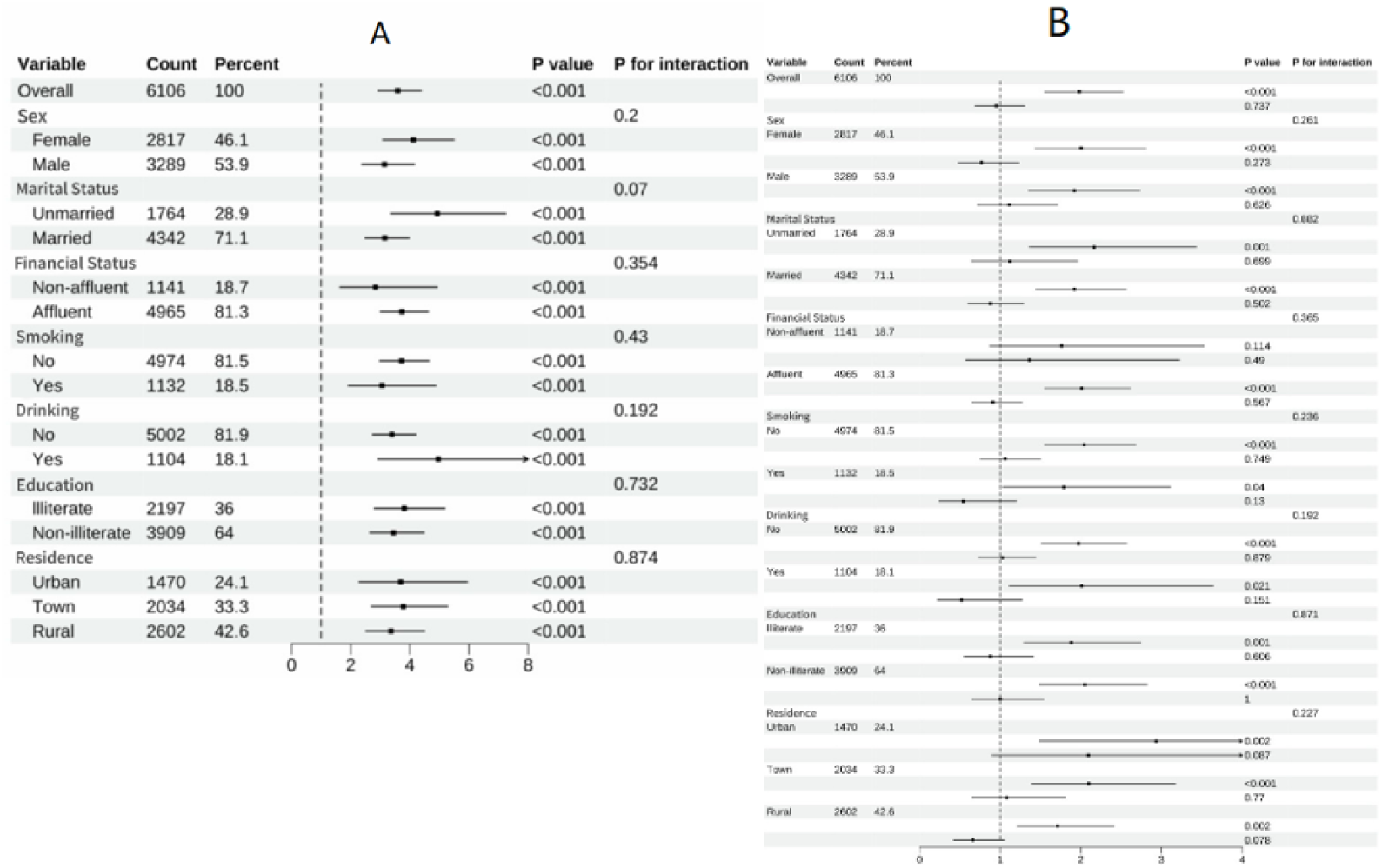
Forest Plot of the Subgroup Analysis for the Association of Sleep Quality and Sleep Duration with Anxiety Mediation Analysis

The results of the mediation analyses are presented in Table 3 and Figure 3. When physical exercise was included as a mediator, the total effect of sleep quality on anxiety was significant (β = 0.110, p < 0.001). The direct effect remained significant (β = 0.108, p < 0.001), accounting for 98.18% of the total effect. A significant indirect effect was observed through physical exercise (β = 0.002, 95% CI: 0.000 to 0.003, p = 0.016), with the bootstrap confidence interval excluding zero. The proportion mediated was 1.56%. Path coefficients indicated that better sleep quality (reverse-coded) was associated with greater physical exercise (β = 0.356, p < 0.001), and that lower levels of physical exercise were associated with higher anxiety (β = 0.235, p = 0.015). As shown in Table 4 and Figure 3, when social participation was tested as a mediator, the total effect of sleep quality on anxiety was significant (β = 0.108, p < 0.001). The direct effect was also significant (β = 0.108, p < 0.001). However, the indirect effect via social participation was not significant (β = 0.000, 95% CI: -0.000 to 0.001, p = 0.410), as the bootstrap confidence interval included zero. Thus, social participation did not demonstrate a significant mediating role in the relationship between sleep quality and anxiety.

**Fig. 3.**
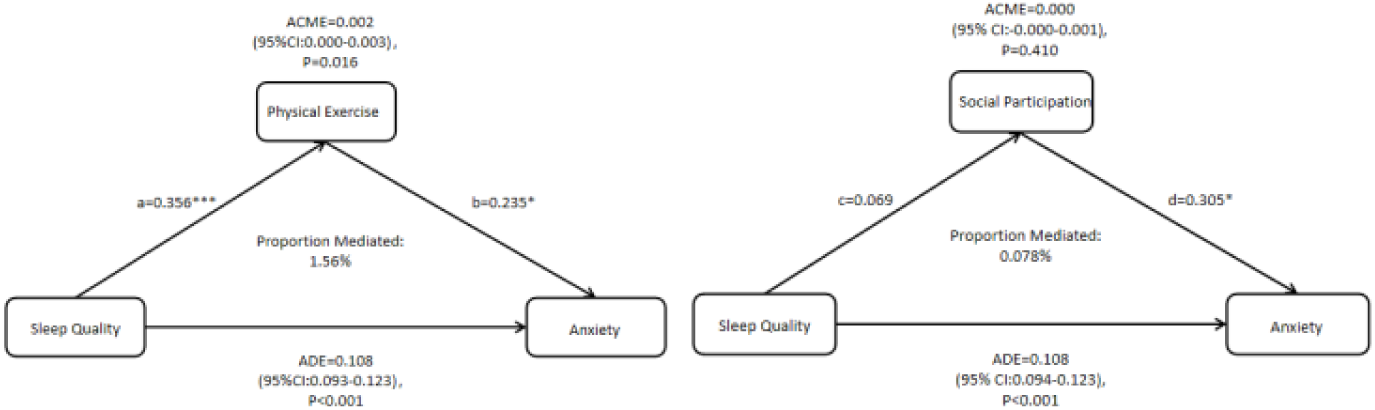
Path Diagram of the Mediating Effects of Physical Exercise and Social Participation Between Sleep Quality and Anxiety

**Table 3.** Mediating Effect of Physical Exercise Between Sleep Quality and Anxiety.

| Pathway | $\beta$ | SE | BootLLCI | BootULCI | p-value |
| --- | --- | --- | --- | --- | --- |
| Total Effect | 0.110 | 0.008 | 0.094 | 0.124 | <0.001 |
| Direct Effect | 0.108 | 0.008 | 0.093 | 0.123 | <0.001 |
| Sleep Quality → Physical Exercise | 0.356 | 0.053 | 0.252 | 0.460 | <0.001 |
| Physical Exercise → Anxiety | 0.235 | 0.096 | 0.047 | 0.423 | 0.015 |
| Sleep Quality → Physical Exercise → Anxiety | 0.002 | 0.001 | 0.000 | 0.003 | 0.016 |

**Table 4.**
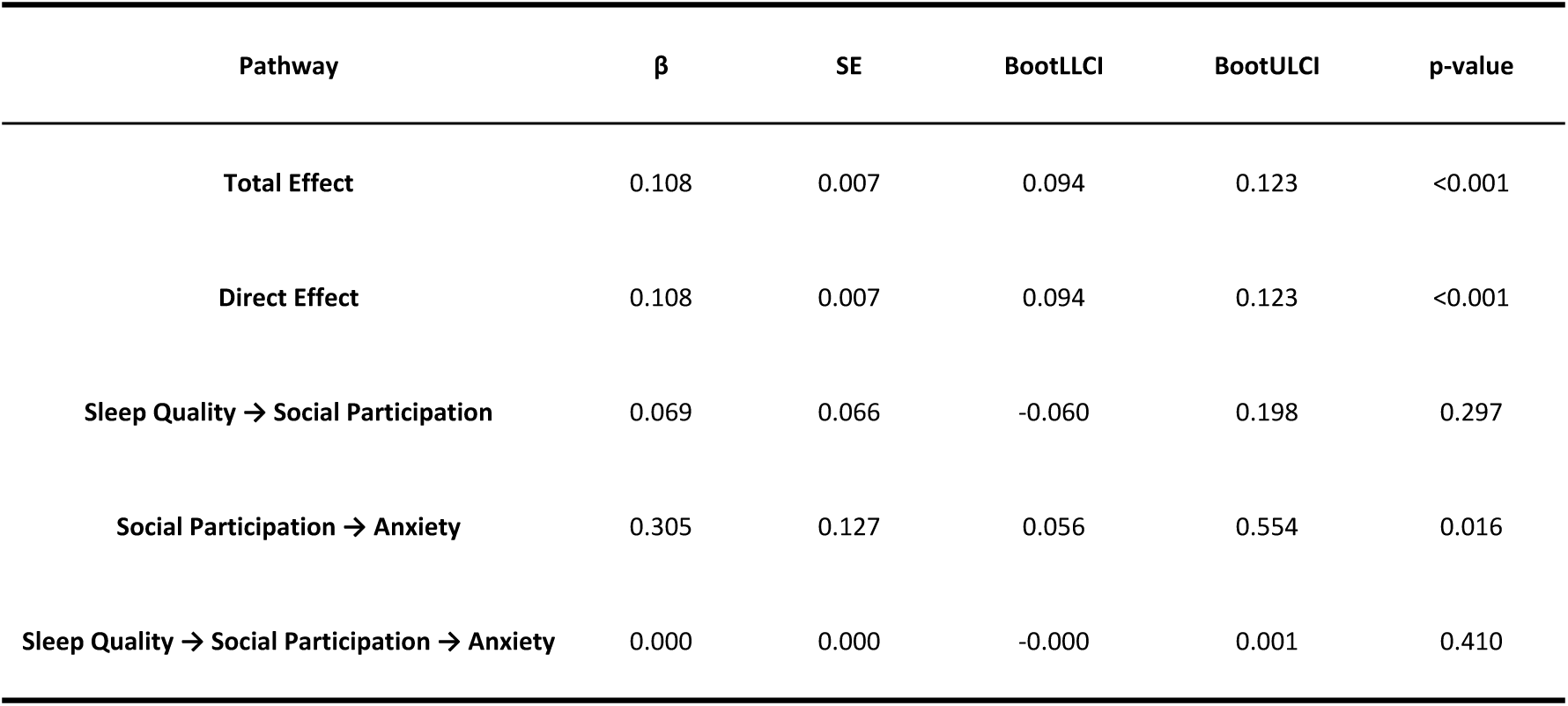
Mediating Effect of Social Participation Between Sleep Quality and Anxiety.

| Pathway | $\beta$ | SE | BootLLCI | BootULCI | p-value |
| --- | --- | --- | --- | --- | --- |
| Total Effect | 0.108 | 0.007 | 0.094 | 0.123 | <0.001 |
| Direct Effect | 0.108 | 0.007 | 0.094 | 0.123 | <0.001 |
| Sleep Quality → Social Participation | 0.069 | 0.066 | -0.060 | 0.198 | 0.297 |
| Social Participation → Anxiety | 0.305 | 0.127 | 0.056 | 0.554 | 0.016 |
| Sleep Quality → Social Participation → Anxiety | 0.000 | 0.000 | -0.000 | 0.001 | 0.410 |

### Mediation Analysis for Sleep Duration

Mediation analyses for short sleep duration are summarized in Table 5 and Figure 4. When physical exercise was examined as a mediator, the total effect of short sleep duration on anxiety was significant (β = 0.063, p < 0.001), as was the direct effect (β = 0.063, p < 0.001). The indirect effect via physical exercise was not statistically significant (β = 0.000, 95% CI: -0.001 to 0.001, p = 0.826), as the bootstrap confidence interval included zero. When social participation was tested as the mediator (Table 6 and Figure 4), the total effect of short sleep duration on anxiety remained significant (β = 0.062, p < 0.001). The direct effect was also significant (β = 0.063, p < 0.001). The indirect effect through social participation was statistically significant (β = −0.001, 95% CI: −0.002 to −0.000, p = 0.008), with the bootstrap confidence interval excluding zero; the estimated proportion mediated was −1.44%. In the path analysis, shorter sleep duration (positively coded) was associated with lower social participation (reverse-coded; β = -0.235, p < 0.001), which in turn was associated with higher anxiety levels (β = 0.334, p = 0.008).

**Fig. 4.**
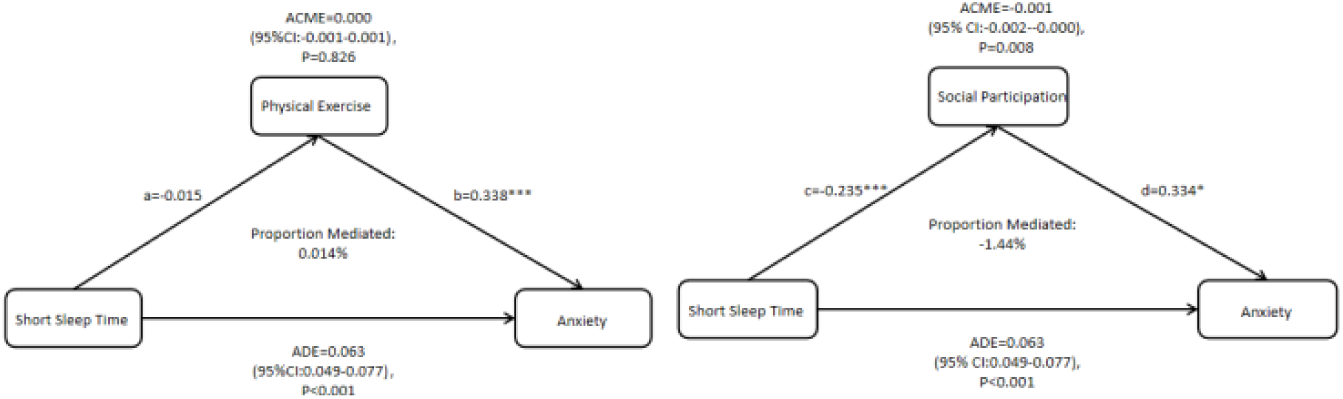
Path Diagram of the Mediating Effects of Physical Exercise and Social Participation Between Short Sleep Duration and Anxiety

**Table 5.** Mediating Effect of Physical Exercise Between Short Sleep Duration and Anxiety.

| Pathway | $\beta$ | SE | BootLLCI | BootULCI | p-value |
| --- | --- | --- | --- | --- | --- |
| Total Effect (c) | 0.063 | 0.007 | 0.049 | 0.077 | <0.001 |
| Direct Effect | 0.063 | 0.007 | 0.049 | 0.077 | <0.001 |
| Short Sleep Time → Physical Exercise | -0.015 | 0.053 | -0.119 | 0.089 | 0.782 |
| Physical Exercise → Anxiety | 0.338 | 0.095 | 0.152 | 0.524 | <0.001 |
| Short Sleep Time → Physical Exercise → Anxiety | 0.000 | 0.001 | -0.001 | 0.001 | 0.826 |

**Table 6.**
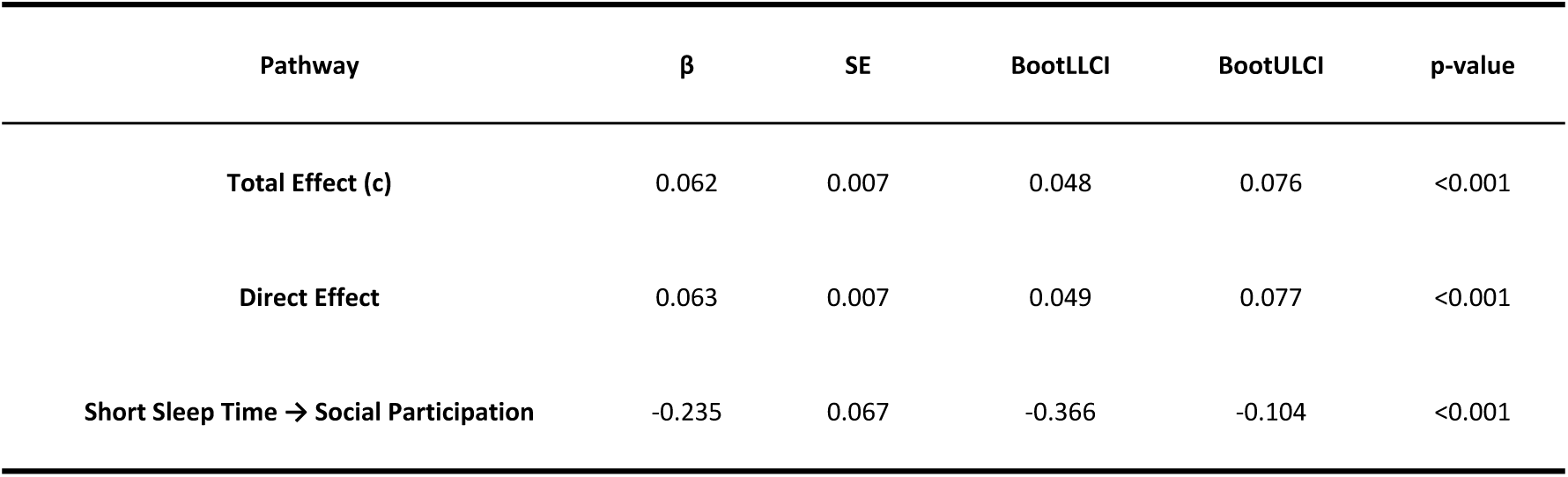
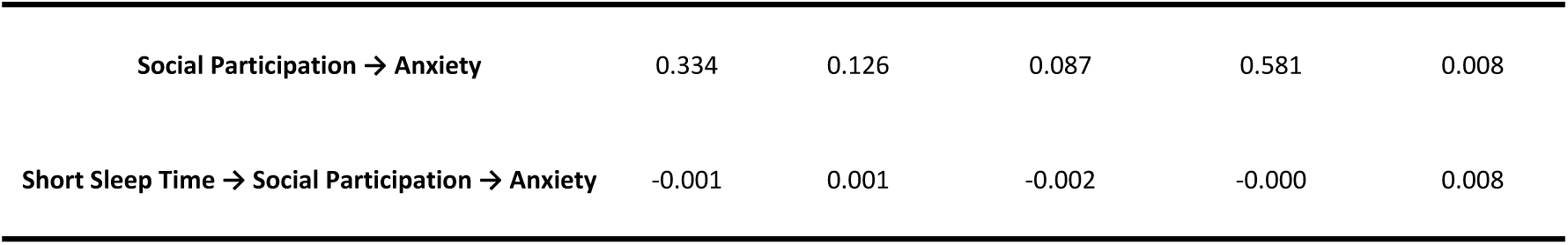
Mediating Effect of Social Participation Between Short Sleep Duration and Anxiety.

### Mediation Analysis for Long Sleep Duration

The mediation analyses for long sleep duration are detailed in Table 7 and Figure 5. When physical exercise was included as a mediator, the total effect of long sleep duration on anxiety was significant (β = -0.046, p < 0.001). The direct effect was also significant (β = -0.047, p < 0.001). The indirect effect through physical exercise was statistically significant (β = 0.001, 95% CI: 0.000–0.002, p = 0.009), with the bootstrap confidence interval excluding zero; the estimated proportion mediated was −2.80%. In the path model, longer sleep duration was associated with lower levels of physical exercise (β = 0.198, p = 0.002), which in turn was associated with higher anxiety (β = 0.349, p < 0.001). When social participation was tested as the mediator (Table 8 and Figure 5), the total effect of long sleep duration on anxiety remained significant (β = -0.045, p < 0.001), as did the direct effect (β = -0.047, p < 0.001). A statistically significant indirect effect via social participation was observed (β = 0.002, 95% CI: 0.001–0.004, p = 0.005), with the bootstrap confidence interval excluding zero; the estimated proportion mediated was −4.78%. Path coefficients indicated that longer sleep duration was associated with lower social participation (β = 0.695, p < 0.001), and lower social participation was associated with higher anxiety levels (β = 0.340, p = 0.007).

**Fig. 5.**
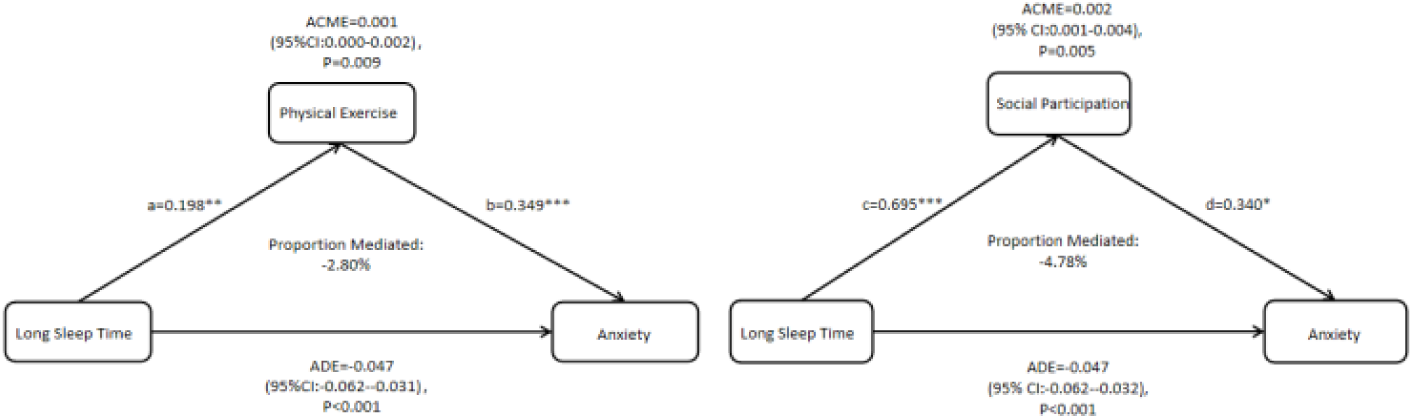
Path Diagram of the Mediating Effects of Physical Exercise and Social Participation Between Long Sleep Duration and Anxiety

**Table 7.** Mediating Effect of Physical Exercise Between Long Sleep Duration and Anxiety.

| Pathway | $\beta$ | SE | BootLLCI | BootULCI | p-value |
| --- | --- | --- | --- | --- | --- |
| Total Effect (c) | -0.046 | 0.008 | -0.061 | -0.030 | <0.001 |
| Direct Effect | -0.047 | 0.008 | -0.062 | -0.031 | <0.001 |
| Long Sleep Time → Physical Exercise | 0.198 | 0.065 | 0.071 | 0.325 | 0.002 |
| Physical Exercise → Anxiety | 0.349 | 0.095 | 0.163 | 0.535 | <0.001 |
| Long Sleep Time → Physical Exercise → Anxiety | 0.001 | 0.001 | 0.000 | 0.002 | 0.009 |

**Table 8.** Mediating Effect of Social Participation Between Long Sleep Duration and Anxiety.

| Pathway | $\beta$ | SE | BootLLCI | BootULCI | p-value |
| --- | --- | --- | --- | --- | --- |
| Total Effect (c) | -0.045 | 0.008 | -0.060 | -0.030 | <0.001 |
| Direct Effect | -0.047 | 0.008 | -0.062 | -0.032 | <0.001 |
| Long Sleep Time → Social Participation | 0.695 | 0.093 | 0.513 | 0.877 | <0.001 |
| Social Participation → Anxiety | 0.340 | 0.126 | 0.093 | 0.587 | 0.007 |
| Long Sleep Time → Social Participation → Anxiety | 0.002 | 0.001 | 0.001 | 0.004 | 0.005 |

## Discussion

This study, involving 6,106 Chinese empty-nest older adults, identified poor sleep quality and short sleep duration as independent associated with anxiety symptoms, with the effect of poor sleep quality being more pronounced. In contrast, the association between long sleep duration and anxiety was not statistically significant, suggesting a minimal role. Subgroup analyses revealed that the associations of both sleep quality and sleep duration with anxiety were consistent across strata defined by sex, marital status, and economic status, with no significant interaction effects. Mediation analyses indicated that physical exercise partially mediated the associations between sleep quality and long sleep duration and anxiety. Social participation also served as a significant mediator in the relationship between sleep duration (both short and long) and anxiety.

Epidemiological evidence and systematic reviews consistently identify sleep disturbances as significant risk factors for anxiety and depression in older adults.^[25]^.For example, a large-sample study in China reported a significant negative correlation between sleep quality and anxiety symptoms, following a dose-response relationship^[26]^. Meta-analytic evidence further suggests that improvements in sleep quality are associated with reductions in anxiety and depression symptoms and enhanced mental well-being.^[27]^.Research involving multi-ethnic populations indicates that while short sleep duration is linked to psychological distress, poor sleep quality demonstrates a stronger association and larger effect size with generalized anxiety disorder and major depression^[28]^.These findings are consistent with the present study, underscoring that sleep quality may play a more pivotal role than sleep duration in the etiology of anxiety.

Sleep may influence anxiety through multiple physiological pathways. Neurological research indicates that sleep disturbances can induce a state of hyperarousal and sleep fragmentation. These changes are accompanied by impaired emotion regulation in the prefrontal cortex, autonomic nervous system (ANS) dysregulation, and activation of the hypothalamic-pituitary-adrenal (HPA) axis and inflammatory responses, collectively exacerbating anxiety symptoms^[29]^. Neuroimaging studies have identified abnormal functional connectivity in emotion-related brain regions among individuals with poor sleep, providing mechanistic insights into the sleep-anxiety link^[30]^. Furthermore, clinical intervention studies suggest that improvements in autonomic function and reductions in inflammation are associated with concurrent improvements in sleep disturbances and anxiety, pointing to shared physiological mechanisms.^[31]^. Together, this evidence supports a complex, bidirectional relationship, wherein sleep disturbances are not merely a consequence but also a significant precipitating and perpetuating factor for anxiety.

This study further identified physical exercise as a significant mediator in the relationships of both sleep quality and long sleep duration with anxiety. This observation is supported by existing literature. For instance, a systematic review and meta-analysis by Solis-Navarro et al^[32]^. demonstrated that regular exercise significantly improves sleep quality in older adults. Similarly, intervention studies, such as that by Goodarzi et al^[33]^. Support the beneficial effects of exercise on anxiety symptoms in this population. Potential mechanisms may include the reduction of inflammation and oxidative stress, enhancement of neuroplasticity, and promotion of endorphin release^[34]^. Therefore, promoting physical exercise represents a promising intervention target for simultaneously improving sleep and reducing anxiety in older adults.

The mediating role of social participation in the sleep-anxiety relationship was relatively circumscribed. Specifically, significant indirect effects were observed only for sleep duration (both short and long) but not for sleep quality. This pattern aligns with existing literature. Liang et al. (2024) reported that while social participation was inversely associated with anxiety and depression, its effect on anxiety was primarily indirect, mediated by reductions in loneliness and depressive symptoms rather than a direct pathway^[35]^. Similarly, Peng et al. (2025) found that among older adults with multimorbidity, the link between social participation and insomnia was fully mediated by frailty, anxiety, and depression, with no direct association remaining after accounting for these factors^[36]^.These findings suggest that the benefits of social participation for anxiety and sleep are largely contingent on its positive effects on mental health and physical function, with limited independent influence.

Several factors may explain the limited or indirect effects observed. The quality of social engagement and the level of emotional support are likely critical, as suggested by studies in Chinese populations highlighting the importance of social support for mental health. Positive effects may be attenuated by poor-quality interactions or interpersonal conflict^[37]^.Furthermore, opportunities for meaningful social participation are often constrained in older adults by declining physical function or mobility limitations, which may further dilute its potential protective role against anxiety and sleep disturbances.

Several limitations of this study should be noted. First, the cross-sectional design hinders causal inference regarding the observed associations. Second, physical exercise and social participation were measured with single items, limiting the assessment of their frequency, intensity, and quality; this may have resulted in an underestimation of their true effects. Third, although multiple covariates were adjusted for, residual confounding by unmeasured factors (e.g., chronic disease burden, medication use) remains possible.

In conclusion, this study demonstrates that poor sleep quality and short sleep duration are independently associated with anxiety among Chinese empty-nest older adults, and identifies physical exercise and social participation as potential mediators. These findings suggest that multi-level interventions are warranted, targeting sleep improvement directly while also promoting physical exercise and social engagement to indirectly mitigate anxiety. Future longitudinal studies incorporating objective sleep measures (e.g., polysomnography) and biomarkers are needed to clarify causality and elucidate the underlying biological and psychosocial mechanisms.

## Conclusion

In conclusion, among Chinese empty-nest older adults, both sleep quality and short sleep duration are independently associated with anxiety symptoms, with sleep quality exerting a stronger association. Physical exercise partially mediates the relationships of both sleep quality and long sleep duration with anxiety, whereas social participation mediates the association specifically between sleep duration and anxiety. These findings underscore the importance of prioritizing sleep quality improvement while concurrently promoting regular physical exercise and social engagement as a multifaceted strategy to mitigate anxiety risk. Future longitudinal research incorporating objective sleep measures is warranted to clarify causal relationships and inform targeted interventions.

## Data availability

This study is based entirely on publicly available secondary data. The Chinese Longitudinal Healthy Longevity Survey (CLHLS) data are open-access and de-identified, and can be obtained from the Center for Healthy Aging and Development Studies (CHADS) at Peking University (https://opendata.pku.edu.cn/dataverse/CHADS).

## Contributions

M.Y.wrote the main manuscript text, data analysis and visualization. M.Y. and Z.Q. enabled conceptualization and provided methodology. J.J. and Z.N. participated in paper review and editing.

